# Neural CXCR4 contributes to neuroimmune modulation of atherosclerosis in a cardio-metabolic *Ldlr^⁻/⁻^* mouse model

**DOI:** 10.64898/2025.12.16.694604

**Authors:** Yaqi Sun, Robby Zachariah Tom, Anna Götz, Sebastian Cucuruz, Annette Feuchtinger, Dominik Lutter, Yvonne Jansen, Christian Weber, Yvonne Döring, Cristina García Cáceres, Aphrodite Kapurniotu, Jürgen Bernhagen, Timo D. Müller, Susanna M. Hofmann

## Abstract

**Background:** Atherosclerosis is a chronic inflammatory disease characterized by metabolic and immune dysregulation. Emerging evidence suggests that specific central nervous system (CNS) regions modulate its progression via neuroimmune cardiovascular interfaces (NICIs). While the C-X-C motif chemokine receptor 4 (CXCR4) is known to participate in atherogenesis by regulating immune cell dynamics and vascular wall responses, the role of neural CXCR4 in atherosclerotic plaque formation remains unclear.

**Methods:** We generated a *Nestin-Cre*–mediated conditional *Cxcr4* knockout mouse model on a low-density lipoprotein receptor–deficient (*Ldlr^⁻/⁻^*) background and fed these mice a western diet (WD) for up to 12 weeks to induce atherosclerosis. To evaluate the impact of neural CXCR4 function, we quantified atherosclerotic plaque burden, systemic metabolic parameters and circulating immune cell profiles, comparing neural *Cxcr4*-deficient mice with corresponding genetic controls. Spatial transcriptomics and RNAscope were employed to map *Cxcr4* mRNA and its non-canonical ligand macrophage migration inhibitory factor (*Mif*) mRNA in neuroimmune-regulatory brain regions, and to assess diet-induced expression changes in relation to neuroinflammatory responses.

**Results:** Neural conditional deletion of *Cxcr4* in *Ldlr*^⁻/⁻^ mice significantly reduced atherosclerotic plaque formation in the aortic arch and aorta, without affecting body weight, lipid levels, glucose tolerance, or circulating immune cells. *Cxcr4* gene expression was found to be uniformly low across hypothalamic subregions implicated in neuroimmune regulation of systemic inflammation and atherogenesis. Importantly, WD feeding did not modify this consistently low expression in male *Ldlr*^⁻/⁻^ mice. In contrast, *Mif* mRNA expression was significantly upregulated in the PVN after 5-day WD feeding, but not after 8 weeks. Exploratory spatial transcriptomic analysis of PVN-containing coronal brain sections from male *Ldlr^⁻/⁻^* mice suggested that 5-day WD exposure is associated with MIF–CXCR4 signaling and downstream neuroinflammatory pathways in the PVN.

**Conclusions:** This study identifies neural CXCR4 as a component of neuroimmune modulation in atherosclerosis, exerting its effect independently of systemic metabolic or inflammatory changes. Short-term WD exposure activated gene expression of the CXCR4 ligand *Mif* in the PVN, pointing to a neuroimmune axis that may promote vascular inflammation and atherosclerotic plaque development. These findings establish a link between CNS CXCR4 and vascular disease and suggest that MIF–CXCR4–dependent neuroimmune pathways may play a role in cardiometabolic risk.

**Trial registration:** Not applicable.

## Background

Atherosclerosis remains a predominant cause of cardiovascular morbidity and mortality worldwide, characterized by chronic inflammation, lipid accumulation, and immune cell infiltration within arterial walls [1, 2]. Although peripheral mechanisms such as dyslipidemia and inflammatory mechanisms underlying atherogenesis have been extensively studied, accumulating evidence increasingly highlights the critical role of the CNS in regulating peripheral metabolic homeostasis, inflammation, and cardiovascular function [3–6]. Recent studies further indicate that specific CNS regions, including the hypothalamus and amygdala, regulate peripheral inflammation and modulate the development and progression of atherosclerosis through NICIs [7]. However, current insights into CNS-mediated regulation of atherogenesis remain limited, particularly regarding the specific neural pathways and molecular signals involved.

CXCR4 is a G-protein-coupled receptor (GPCR) widely expressed in various tissues and cells, playing essential roles in cell migration, immune responses, and inflammatory processes [8, 9]. CXCR4 signaling is well known to play an important role in atherosclerosis; however, accumulating evidence indicates that its effects are highly context- and cell type–dependent, with both proatherogenic and atheroprotective functions [10]. Activation of CXCR4 by its non-cognate ligand MIF facilitates recruitment of atherogenic leukocytes and exacerbates inflammation within atherosclerotic lesions, and selective pharmacological blockade of the MIF–CXCR4 axis reduces arterial leukocyte adhesion and attenuates atherosclerosis in hyperlipidemic apolipoprotein E–deficient (*Apoe*^⁻/⁻^) mice [11, 12]. In contrast, CXCR4 can exert atheroprotective effects in specific vascular cell types. Endothelial CXCR4 signaling preserves vascular integrity and limits leukocyte adhesion by maintaining endothelial barrier function, while smooth muscle cell CXCR4 promotes a contractile phenotype and stabilizes plaques [13, 14].

In the CNS, CXCR4 is constitutively expressed in various cell types including neurons, microglia, astrocytes, and oligodendrocytes, across multiple brain regions such as the cortex, hippocampus, and hypothalamus [15–17]. Neural CXCR4 signaling is involved in neuroinflammation, immune cell trafficking, and neuroendocrine modulation [18–21]. However, whether neural CXCR4 signaling directly impacts on atherosclerotic lesion development remains unexplored.

In this study, we examined the role of neural CXCR4 in cardiometabolic disease using a *Nestin-Cre*–mediated conditional *Cxcr4* knockout on an *Ldlr*⁻/⁻ background. Employing spatial transcriptomics and RNAscope, we mapped *Cxcr4* and its ligand *Mif* within neuroimmune-regulatory regions and determined whether Western Diet (WD) exposure alters their gene expression in patterns consistent with the initiation of neuroimmune signaling linked to vascular inflammation and atherosclerotic progression.

## Materials and methods

### Animals and diet

All mice were on a *C57BL/6J* background. *Ldlr*^⁻/⁻^ mice and *Nestin-Cre* mice were purchased from Jackson Laboratories (https://www.jax.org/strain/002207; ME, USA). The *Cxcr4-floxed* (*Cxcr4^fl/fl^)* mice, kindly provided by Dr. Y. Zou (Columbia University, New York, US), were backcrossed into a C57BL/6 background and crossed with C57BL/6 *Ldlr*^⁻/⁻^ mice. To study neural CXCR4 function in atherosclerosis, *Cxcr4^fl/fl^Ldlr*^⁻/⁻^mice were crossbred with *Nestin-Cre* mice, resulting in *Nestin-Cre^+^Cxcr4^fl/fl^Ldlr*^⁻/⁻^ mice, in which *Cxcr4* was specifically deleted in *Nestin*-positive neural cells on an *Ldlr*^⁻/⁻^ background (Fig. 1A). Littermate *Nestin-Cre^-^Cxcr4^fl/fl^Ldlr*^⁻/⁻^ mice served as controls. For atherosclerosis studies, 12-week-old male and female *Nestin-Cre^+^Cxcr4^fl/fl^Ldlr*^⁻/⁻^ mice and control littermates were fed a WD containing 21% fat and 0.15-0.2% cholesterol (Sniff TD88137) for 12 weeks. To assess diet-induced changes in *Cxcr4* and *Mif* mRNA expression, male and female *Ldlr^⁻/⁻^* mice were fed either a standard diet (SD; Altromin 1318) or a WD (Sniff TD88137) for either 5 days or 8 weeks.

**Figure 1.**
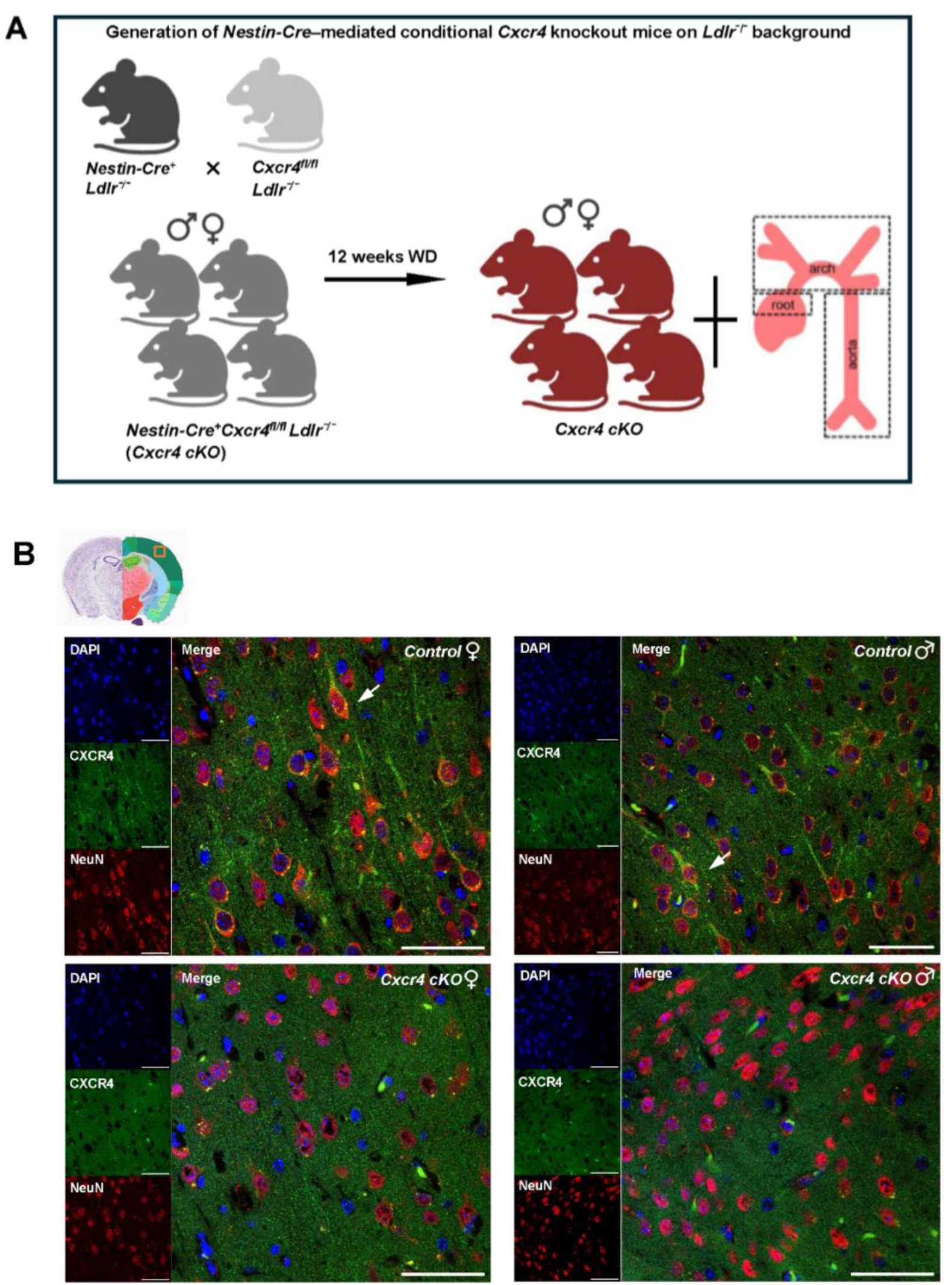
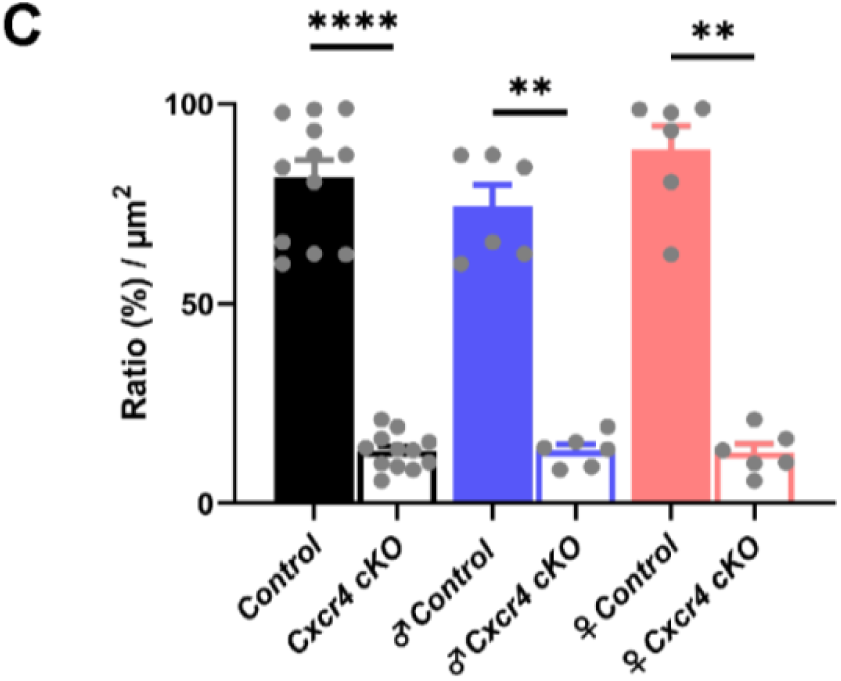
Establishment of *Nestin-Cre*–mediated conditional *Cxcr4* knockout mice on *Ldlr*^⁻/⁻^ background. (A) Experimental schematic illustrating the generation of *Nestin-Cre^+^Cxcr4^fl/fl^Ldlr*⁻^/^⁻ (*Cxcr4 cKO*) mice. Both male (♂) and female (♀) *Cxcr4 cKO* mice and their littermate *Nestin-Cre^-^Cxcr4^fl/fl^Ldlr*^⁻/⁻^ (*Control*) mice were fed WD for 12 weeks prior to tissue collection for analysis. (B) Immunofluorescence staining of CXCR4 (green), neuronal marker NeuN (red), and nuclear marker DAPI (blue) in cortical sections from male (♂) and female (♀) *Cxcr4 cKO* mice and their littermate *Control* mice. Smaller panels showed individual staining for CXCR4, NeuN, and DAPI. White arrows indicate co-localization of CXCR4 with neurons. The orange box in the schematic mouse brain atlas indicates the anatomical location of the imaged cortical region. Scale bar, 50 µm. (C) Quantification of cortical CXCR4-positive neurons in both male (♂) and female (♀) *Cxcr4 cKO* mice and *Control* mice. Data represents the ratio of CXCR4-positive cortical neurons to the total number of cortical neurons. Results are presented as mean ± SEM (n = 6 mice/group). Statistical significance was determined by Mann–Whitney U tests (****, *p* < 0.0001; **, *p* < 0.01).

All mice were double-housed and maintained at 22+/-2 °C, 55 +/- 10% relative humidity, and a 12-h light/dark cycle with free access to food and water. Body fat mass was measured in conscious mice using 1H magnetic resonance spectroscopy (EchoMRI-100; Echo Medical Systems). All procedures were approved by the local Animal Use and Care Committee and the local authorities of Upper Bavaria, Germany in accordance with European and German animal welfare regulations.

### Histology

For analysis of mouse atherosclerotic lesions, the aortic root and thoraco-abdominal aorta were stained for lipid depositions with Oil-Red-O. In brief, hearts with aortic root were embedded in Tissue-Tek O.C.T. compound (Sakura Finetek, Torrance, CA, USA) for cryo-sectioning. Atherosclerotic lesions were quantified in 5 μm transverse sections and averages were calculated from 3-5 sections. The aorta was opened longitudinally, mounted on glass slides and *en face*- stained with Oil-Red-O. Aortic arches with main branch points (brachiocephalic artery, left subclavian artery and left common carotid artery) were fixed with 4% paraformaldehyde and embedded in paraffin. Lesion size was quantified after HE-staining of 3 transverse sections.

### Biochemical analysis

Tail blood glucose levels (milligrams per deciliter) were measured with a TheraSense Freestyle glucometer (Abbott Diabetes Care, Inc., Alameda, CA). Mice were euthanized using CO₂. Blood was collected from the inferior vena cava, mixed with EDTA, and immediately placed on ice. Plasma was separated by centrifugation at 4800 g for 10 min at 4 °C. Plasma levels of insulin (Crystal Chem, Elk Grove Village, IL, USA), total cholesterol (Thermo Fisher Scientific, Waltham, MA, USA), and triglycerides (Wako Chemicals, Neuss, Germany) were measured according to the manufacturers’ instructions. For lipoprotein separation, samples from each treatment group were pooled and analyzed in a fast-performance liquid chromatography (FPLC) with two Superose 6 columns connected in series as described previously [22].

### Immunofluorescent (IF) staining

Mice were euthanized with CO₂ and transcardially perfused with 0.9% saline followed by 4% paraformaldehyde (PFA) in phosphate-buffered saline (PBS, pH 7.4). Brains were then extracted, post-fixed in 4%PFA at 4°C for 48 h and then transferred to 30% sucrose in Tris-buffered saline (TBS; pH 7.2) for at least 48 hours until they sank to the bottom of the container. Brain samples were acclimated to the cryostat (Leica Biosystems, Nussloch, Germany) at −20 °C for 30 min and then sectioned coronally at 30 µm thickness. Subsequently, the brain slices were collected and rinsed in 0.1 M TBS. The collected brain slices were permeabilized with 0.5% Triton X-100 in TBS for 30 min followed by washing in TBS. For CXCR4 staining, brain sections were treated with λ protein phosphatase (800 U; Sigma-Aldrich, St. Louis, MO, USA) in Protein MetalloPhosphatases buffer with MnCl_2_ provided by manufacturer for 30 min at room temperature. Subsequently brain sections were blocked in 5% BSA and 0.5% Triton X-100 in TBS for 1 h and incubated overnight at 4°C with primary antibody (rabbit anti-CXCR4, 1:500; Abcam, Cambridge, UK; cat. no. ab181020) with mouse anti-NeuN (1:500; Abcam; cat. no. ab104224). Sections were washed in TBS, incubated with secondary antibodies; Alexa Fluor 555 donkey anti-rabbit (1:1000; Abcam; cat. no. ab150062) or Alexa Fluor 647 donkey anti-mouse (1:1000; Abcam; cat. no. ab150111) for 2 h at room temperature and stained with 4′,6-diamidino-2-phenylindole (DAPI) solution (1:1000; Thermo Fisher Scientific; cat. no. 62248) for 5 min. After a final wash series with TBS, sections were mounted on gelatin pre-coated glass slides with ProLong Diamond Antifade Mountant (Thermo Fisher Scientific; cat. no. P36961), and a coverslip was mounted onto the sections for image quantification. Confocal pictures were taken with Leica SP5 scanning confocal microscope.

### RNAscope

Mice were euthanized with CO₂ and then subsequently perfused transcardially with 0.9% saline followed by 4% paraformaldehyde (PFA) in phosphate-buffered saline (PBS, pH 7.4). Brains were then extracted and kept in 4% PFA solution at 4°C overnight, and transferred first to 10%, then 20%, and finally 30% sucrose in 0.1 M TBS solution each consecutive day. The brains were acclimated to cryostat (Leica Biosystems) temperature of -20°C for a period of 1 – 2 h and then sliced at 10 μm sections. The slices were immediately mounted onto SuperFrost Plus Gold slides (Fisher Scientific) and kept in -80°C until further use. Slides were processed within one month. All staining procedures were performed using the RNAscope Fluorescent Multiplex Kit (Advanced Cell Diagnostics, Newark, CA, USA; cat. no. 323100) following the manufacturer’s instructions. Briefly, slides were warmed up for 30 min at 60°C, then incubated in 4% PFA solution for 15 min. Next, slides were treated with sequential 5 min-long steps of dehydration with 50%, 70%, and 100% ethanol. Sections were dried at 60°C for 30 minutes, dehydrated with an increasing ethanol gradient, pre-treated with hydrogen peroxide (ACD; cat. no. 322381) and boiled in target retrieval reagent (ACD; cat. no. 322000). After dehydrating in pure ethanol, sections were surrounded by a hydrophobic barrier drawn with an ImmEdge Hydrophobic Barrier Pen (Vector Laboratories, Newark, CA, USA; cat. no. H-4000) and left to dry overnight. The following day, slides were incubated with Protease III (ACD; cat. no. 322340) for 30 min at 40 °C, then hybridized with target probes Mm-*Cxcr4* (ACD; cat. no. 400451) and Mm-*Mif* (ACD; cat. no. 432471) for 2 h at 40°C in a hybridization (HybEZ™) oven. Signal amplification was achieved using amplifiers AMP1-3 and an Opal reagent (Opal 570, PerkinElmer, Waltham, MA, USA; cat. no. FP1488001KT, and Opal 690, PerkinElmer; cat. no. FP1497001KT). Sections were counterstained with DAPI (provided in kit) for 30 seconds and mounted with ProLong Diamond Antifade Mountant (Thermo Fisher Scientific, cat. no. P36961). A Zeiss Axioscan 7 digital slide scanner (Carl Zeiss Microscopy GmbH, Oberkochen, Germany) equipped with a Plan-Apochromat 20×/0.8 objective and ZEN 3.1.1 (blue edition) software was used for image acquisition, and the final images were generated by maximum-intensity projection of z-stacks using Zeiss ZEN 3.1.1 (blue edition) software. The resulting RNAscope images were subsequently processed and analyzed using QuPath v0.4.2 (University of Edinburgh, Edinburgh, UK). Anatomical boundaries for hypothalamic and amygdala subregions were delineated according to the Allen Mouse Brain Atlas (https://atlas.brain-map.org/atlas) and applied to the images prior to quantification. Nuclear boundaries were defined based on DAPI counterstaining, and automatic nuclei segmentation was performed using the QuPath cell detection algorithm. Within each segmented nucleus, *Mif* and *Cxcr4* mRNA puncta were detected as fluorescent dots. For each region, the mean number of dots per nuclei was calculated as a measure of transcript abundance.

### Genome-wide FFPE spatial expression profiling

#### Preparation of FFPE samples

Brain samples from male *Ldlr*^⁻/⁻^ mice on 5-day SD or WD feeding were collected and formalin treated, embedded in Paraffin. The sectioning was performed using a microtome (Thermo Fisher Scientific, model HM340E), utilizing a fresh blade and ensuring thorough decontamination with RNAse AWAY (Thermo Fisher Scientific; cat. no. 7002). The quality of preserved RNA in the FFPE brain blocks was evaluated based on the percentage of RNA fragments above 200 base pairs (DV200).

For each FFPE tissue block, three consecutive 10 μm sections were collected, and RNA was purified using the RNeasy FFPE Kit (Qiagen, Hilden, Germany; cat. no. 73504). The purified RNA was then analyzed on an Agilent 2100 Bioanalyzer and samples showing DV200 values above 50% were selected for further processing. Microtome sections (thickness 5 µm) were placed on microscope slides (with allowable area on the back of the slides) after floating on a water bath at 43°C. After sectioning, the slides were dried vertically at 42°C in a hybridization oven (HB-1000 Hybridizer, LabRepCo, Horsham, PA, USA) for 3 h. The slides were then placed inside a slide mailer in a desiccator and kept overnight at room temperature to ensure proper drying. After overnight drying, deparaffinization and decrosslinking were performed according to the *Visium CytAssist for FFPE – Deparaffinization, Decrosslinking, IF Staining & Imaging* protocol (CG000519; 10x Genomics, Pleasanton, CA, USA).

#### IF Staining & Imaging

IF staining and imaging were performed according to the *Visium CytAssist for FFPE – Deparaffinization, Decrosslinking, IF Staining & Imaging* protocol (CG000519; 10x Genomics). In brief, brain sections were incubated in 1x blocking buffer for 30 min at room temperature followed by incubation with a primary antibody solution Mouse anti-NeuN (1: 50; Abcam; cat. no. ab104224) for 1 h at room temperature in the dark. Sections were washed three times, incubated in secondary antibody solution with Alexa Fluor 555 donkey anti-rabbit (1:200; Abcam; cat. no. ab150062), Alexa Fluor 647 donkey anti-mouse (1:200; Abcam; cat. no. ab150111) and DAPI solution (1:100; Thermo Fisher Scientific; cat. no. 62248) for 20 min at room temperature in the dark. After three washes, sections were treated with TrueBlack® Lipofuscin Autofluorescence Quencher (Biotium, Fremont, CA, USA; cat. no. 23007) at room temperature in the dark for 30 sec. After that, a coverslip was mounted onto the sections for image quantification. IF images were acquired with a Zeiss Axioscan 7 Microscope digital slide scanner (Carl Zeiss Microscopy GmbH) equipped with a Plan-Apochromat 20×/0.8 objective and ZEN 3.1.1 (blue edition) software.

#### Visium spatial transcriptomics and sequencing

All spatial transcriptomics procedures were performed according to the *Visium CytAssist Spatial Gene Expression User Guide* (CG000495; 10x Genomics). The protective coverslip placed on the standard glass slides was carefully removed prior to probe hybridization. Following the probe hybridization and probe ligation steps, two standard glass slides and one Visium CytAssist Gene Expression slide (with two 11 × 11 mm capture areas) were placed in the CytAssist instrument, ensuring that the tissue sections on the standard slides were precisely aligned with the two Visium Capture Areas. Within the instrument, a brightfield image was captured to provide spatial orientation for data analysis, followed by permeabilization of the tissue and transfer of transcriptomic probes to the Visium slide. After the probes were extended, the samples were eluted and transferred to new tubes to initiate the process of constructing gene expression libraries. The final gene expression libraries were sequenced on a NovaSeq 6000 platform (Illumina, San Diego, CA, USA) at a minimum depth of 50,000 read pairs per capture spot covered by tissue. Sequencing libraries were demultiplexed using bcl2fastq software (Illumina). The resulting FASTQ files were processed with the Space Ranger pipeline (v2022.0705.1; 10x Genomics) using the GRCh38-2020-A reference genome.

#### Data processing

For each sample (Sample_SD and Sample_WD), filtered feature–barcode matrices (SpaceRanger v1.3.2) were imported into Seurat (v4.0.1). Low quality spots with fewer than 500 UMIs or fewer than 300 detected genes were excluded [23]. Each sample matrices were normalized with SCTransform [24]. The resulting Seurat objects were then merged into a single object while retaining sample metadata (diet group, section ID). Dimensionality reduction was performed on the merged object using principal component analysis (PCA). The top 30 principal components were used to construct a shared nearest-neighbor (SNN) graph with FindNeighbors, followed by unsupervised Louvain clustering using FindClusters (resolution = 1.0). For visualization, clusters in UMAP space were visualized with the DimPlot function, and spatially resolved distributions of clusters were overlaid on the IF images with the SpatialDimPlot function. To identify the PVN-enriched cluster (workflow showed in Supplementary Figure A), Cluster 1 that spatially encompassed and extended beyond the anatomical PVN region was extracted. A total of 725 spatial spots from Cluster 1 (357 from Sample_SD and 368 from Sample_WD) were re-analyzed by PCA (top 30 PCs) for dimensionality reduction, followed by graph-based reclustering using the Louvain algorithm with a resolution of 0.5. Spatial distributions of the resulting six subclusters were visualized on the IF images using the SpatialDimPlot function. Marker gene analysis across the six subclusters within Cluster 1 (725 spots pooled from both diet groups) was performed using Seurat’s FindAllMarkers function (Wilcoxon rank-sum test) after PrepSCTFindMarkers, with parameters set to only.pos = FALSE, min.pct = 0.25, and logfc.threshold = 0.25. Heatmaps of the top five marker genes per subcluster were visualized using the pheatmap R package. Spatial plot of *Mif*, *Cxcr4*, and *Cxcl12* between SD and WD groups in the PVN-enriched cluster were visualized using Seurat’s SpatialFeaturePlot. Differential expression of *Mif*, *Cxcr4*, and *Cxcl12* between SD and WD groups within the PVN-enriched cluster was quantified using normalized expression values from individual spatial spots. Comparisons were performed with a two-sided Wilcoxon rank-sum test. Results were visualized as boxplots, where boxes indicate the median and interquartile range and dots represent individual spot-level values. Statistical significance was defined as *p* < 0.05. Differentially expressed genes (DEGs) between SD and WD groups within the PVN-enriched cluster were identified using a Wilcoxon rank-sum test implemented in Seurat’s FindMarkers function, with thresholds set at |log₂FC| > 1 and FDR < 0.05. The resulting list of DEGs was uploaded to Ingenuity Pathway Analysis (IPA; QIAGEN, Redwood City, CA, USA; https://www.qiagenbioinformatics.com/products/ingenuity-pathway-analysis) for Core Analysis. Canonical pathways, upstream regulators, and disease/function annotations were generated, with pathways considered significant at –log_10_(*p*)> 1.3 (equivalent to p < 0.05). IPA activation Z-scores were used to predict pathway activity, with Z ≥ 2 indicating activation and Z ≤ –2 indicating inhibition. To specifically characterize CXCR4 signaling pathway, genes annotated by IPA as members of the canonical CXCR4 signaling pathway were extracted and visualized in R using ggplot2. Genes were grouped into functional modules, including G-protein signaling, Rho-GTPase, adenylyl cyclase, PI3K–AKT pathway, MAPK pathway, cell migration, transcription factors, PLC–PKC pathway, ligand/receptor, and others.

### Statistical analysis

All statistical analyses were performed using GraphPad Prism (v8.2.1) and R (v4.1.0). Data are presented as mean ± SEM unless otherwise indicated. Comparisons between two groups were conducted using Welch’s t-test or Mann–Whitney U test, depending on data distribution. For spatial transcriptomics analyses, DEGs between SD and WD conditions were assessed using the Wilcoxon rank-sum test. Statistical significance was defined as p ≤ 0.05.

## Results

### Generation of *Nestin-Cre*–mediated conditional *Cxcr4* knockout mice on *Ldlr*^⁻/⁻^ background

To investigate the specific role of CXCR4 in *Nestin*-expressing neural cells during atherogenesis, male and female *Nestin-Cre^+^Cxcr4^fl/fl^Ldlr*^⁻/⁻^ (*Cxcr4 cKO*) mice and their littermate *Nestin-Cre^-^Cxcr4^fl/fl^Ldlr*^⁻/⁻^ (*Control*) mice were fed an atherogenic WD for 12 weeks. Atherosclerotic lesion formation was analyzed at the aortic arch, aortic root and aorta (Figure 1A). To verify successful neural-specific deletion of *Cxcr4*, immunofluorescence staining was performed on brain sections. Neuronal CXCR4 expression in the cortex was significantly reduced by 71.4% in males and 88.5% in females compared to controls (Figure 1B). These results demonstrate that our *Nestin-Cre*–mediated conditional *Cxcr4* deletion provides a functional knockdown model for the exploration of the biological role of neural CXCR4 in atherosclerosis.

### Neural-specific knockout of *Cxcr4* reduces atherosclerotic plaque formation in the aortic arch and aorta

After 12 weeks of WD feeding, *Cxcr4 cKO* mice exhibited significantly reduced atherosclerotic lesions in the aortic arch, as shown by representative images (Figure 2A) and confirmed by quantitative analysis (Figure 2C). When male (♂) and female (♀) mice were analyzed together, lesion area in the arch was significantly decreased in *Cxcr4 cKO* mice. When analyzed separately by sex, both male (♂) and female (♀) *Cxcr4 cKO* mice displayed a consistent trend toward reduced plaque burden, although these differences did not reach statistical significance within each sex (Figure 2C).

**Figure 2.**
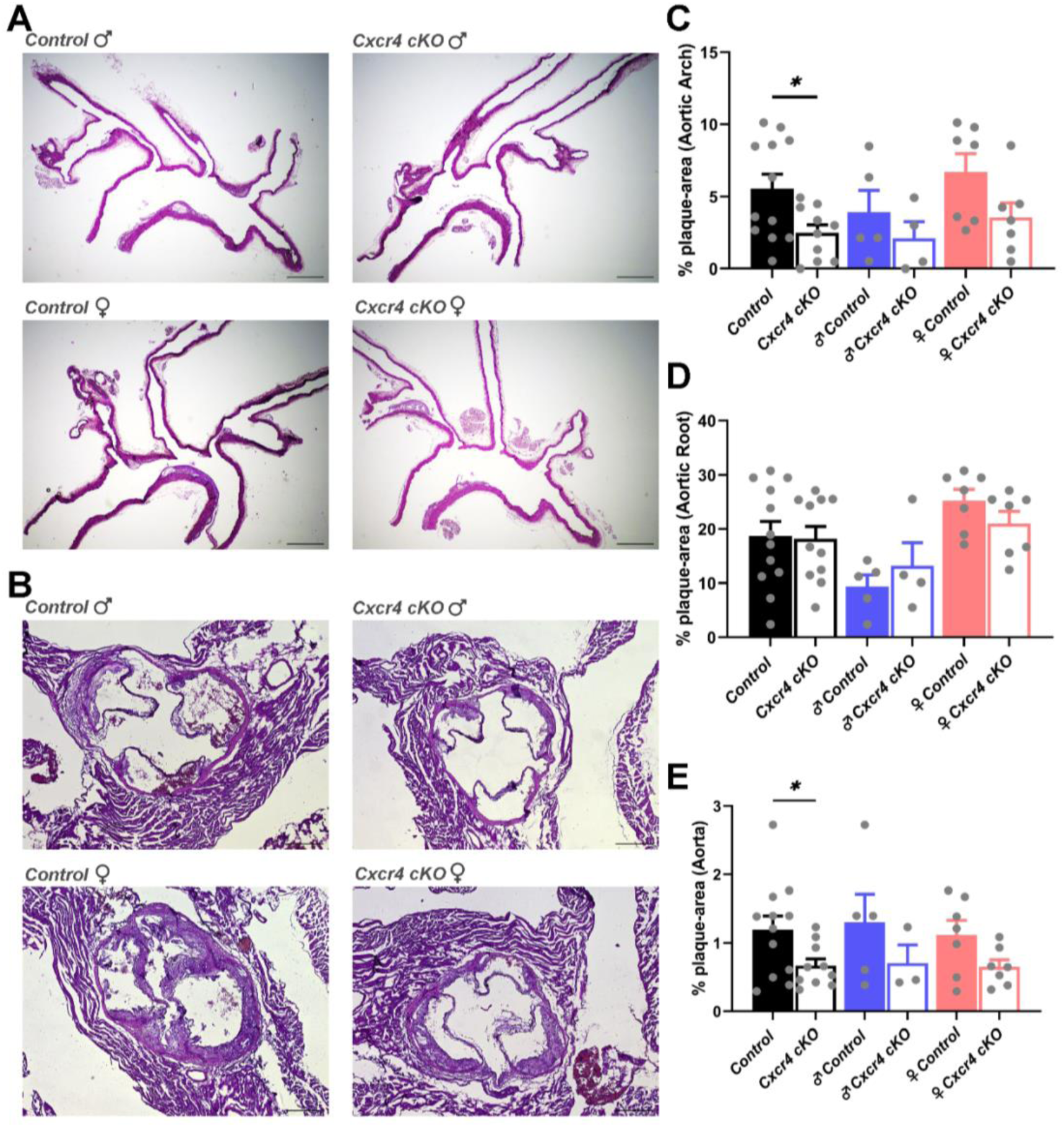
Neural-specific conditional knockout of *Cxcr4* reduces atherosclerotic plaque formation in the aortic arch and aorta. (A, B) Representative images of atherosclerotic lesions in the aortic arch (A) and aortic root (B) from male (♂) and female (♀) *Control* and *Cxcr4 cKO* mice. The aortic arch was analyzed using en face Oil Red O staining, and the aortic root was assessed using H&E-stained cross-sections. Scale bar, 50 µm. (C, D, E) Quantification of atherosclerotic plaque area in the aortic arch (C), aortic root (D), and whole aorta (E). Bar graphs show plaque area as a percentage of total vessel area (mean ± SEM) for combined male (♂) and female (♀) mice (black bars), males (♂) only (blue bars), and females (♀) only (red bars). Statistical significance was determined by Welch’s *t*-test (*, *p* < 0.05).

A similar pattern was observed in the whole aorta, where combined analysis revealed a significant reduction in plaque area in *Cxcr4 cKO* mice, while separate male (♂) and female (♀) evaluations again demonstrated parallel downward trends without statistical significance (Figure 2E).

In contrast, atherosclerotic lesion size at the aortic root was comparable between genotypes, with no detectable differences in either combined or sex-separated analyses (Figure 2B, D). These findings indicate that neural *Cxcr4* deletion exerts a region-specific protective effect, preferentially limiting atherosclerosis in the aortic arch and whole aorta, but not in the aortic root.

### Neural-specific deletion of *Cxcr4* does not alter WD-induced metabolic disease

To determine whether the atheroprotective effects of neural-specific *Cxcr4* deletion were mediated by systemic metabolic changes, we assessed major metabolic parameters in male (♂) and female (♀) *Cxcr4 cKO* and *Control* mice after 12 weeks of WD feeding. These analyses included body weight gain (%), body composition (fat mass, expressed as % body weight), and plasma levels of insulin, triglycerides, total cholesterol, and cholesterol distribution across VLDL, LDL, and HDL fractions. No significant differences were detected between *Control* and *Cxcr4 cKO* mice in any of these metabolic parameters, either when males (♂) and females (♀) were analyzed together or when examined separately by sex (Figure 3A-F). The observed higher cholesterol levels in the VLDL fraction in female mice compared to males are consistent with previously reported sex-related differences in lipid metabolism [25]. These results suggest that neural-specific *Cxcr4* deletion exerts its atheroprotective effects through mechanisms independent of systemic metabolic alterations.

**Figure 3.**
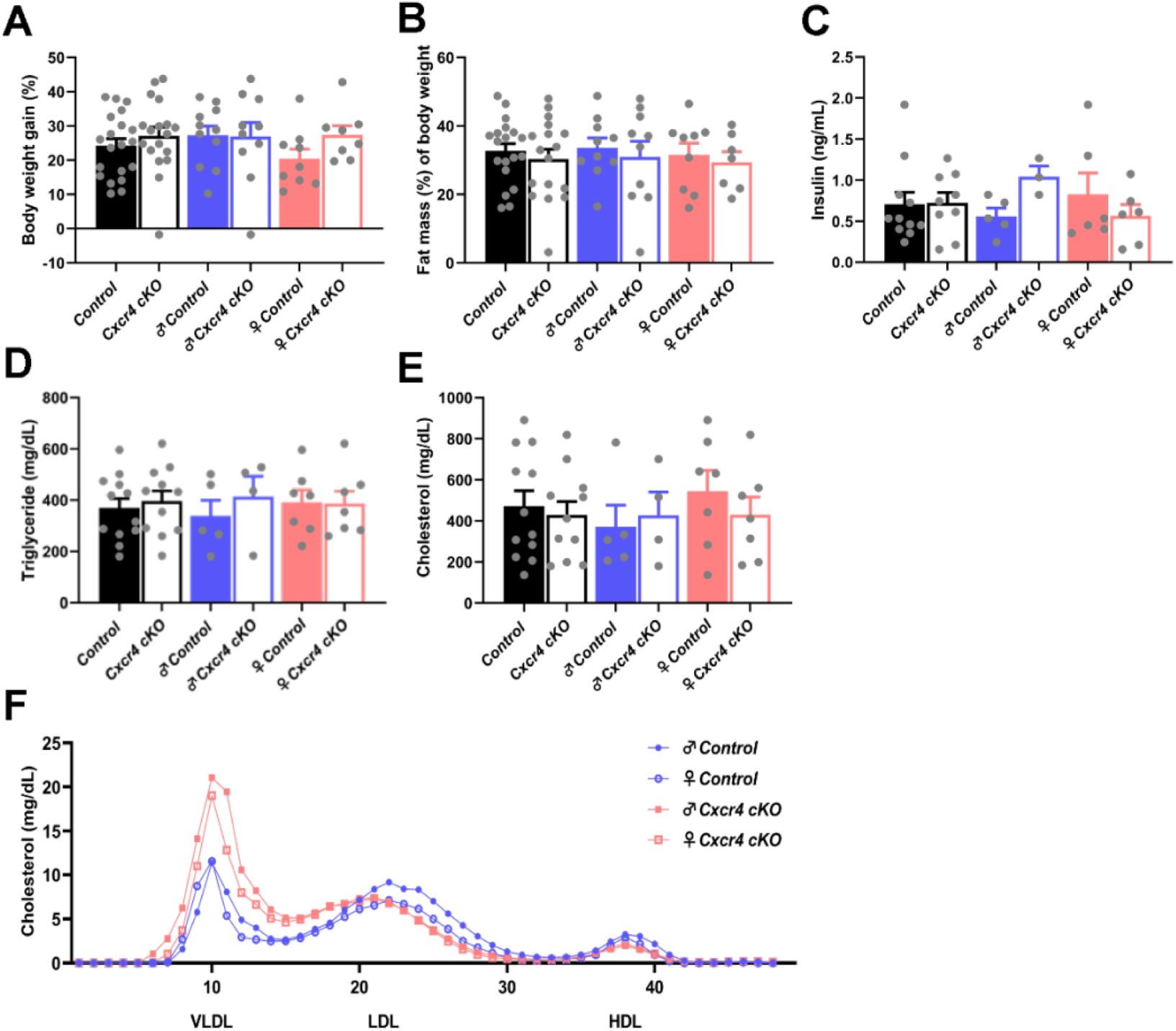
Systemic metabolic parameters in male (♂) and female (♀) *Cxcr4 cKO* mice and *Control* mice after 12 weeks of WD feeding. (A) Body weight gain (%), Body weight gain (%) was calculated as (final body weight – initial body weight) / initial body weight × 100. (n=7-10 per group). (B) Fat mass (% of body weight; n=7-10 per group). (C) Plasma total triglyceride (mg/dL; n=4-7 per group). (D) Plasma total cholesterol (mg/dL, n=4-7 per group). (E) Plasma cholesterol distribution in VLDL, LDL, and HDL fractions determined by FPLC in pooled plasma from male (♂) and female (♀) *Cxcr4 cKO* mice and *Control* mice (n = 4–7 per group). (F) Plasma insulin (ng/mL; n=4-6 per group).

### WD feeding does not modify the *Cxcr4* gene expression in hypothalamic and amygdala subregions of male *Ldlr⁻^/^⁻* mice

To identify specific brain regions involved in CXCR4-mediated modulation of atherogenesis, we first performed RNAscope *in situ* hybridization targeting *Cxcr4* mRNA in *Ldlr^⁻/⁻^* mice, focusing on CNS regions implicated in neuroimmune regulation of peripheral inflammation and atherosclerosis, including hypothalamic subregions (PVN, dorsomedial hypothalamus [DMH], ventromedial hypothalamus [VMH], arcuate nucleus [Arc], and lateral hypothalamic area [LHA]) and amygdala subregions (basolateral amygdala, anterior part [BLAa], and central amygdala [CeA]) [7]. In preliminary experiments, male and female *Cxcr4 cKO* and *Control* mice showed comparable atherosclerotic plaque burden as well as similar metabolic responses to WD feeding, with no detectable sex-dependent differences. To reduce variability and overall animal use, RNAscope were therefore conducted in male *Ldlr^⁻/⁻^* mice. RNAscope detected *Cxcr4* mRNA signals in these regions, though the expression intensity was low (Figure 4A).

**Figure 4.**
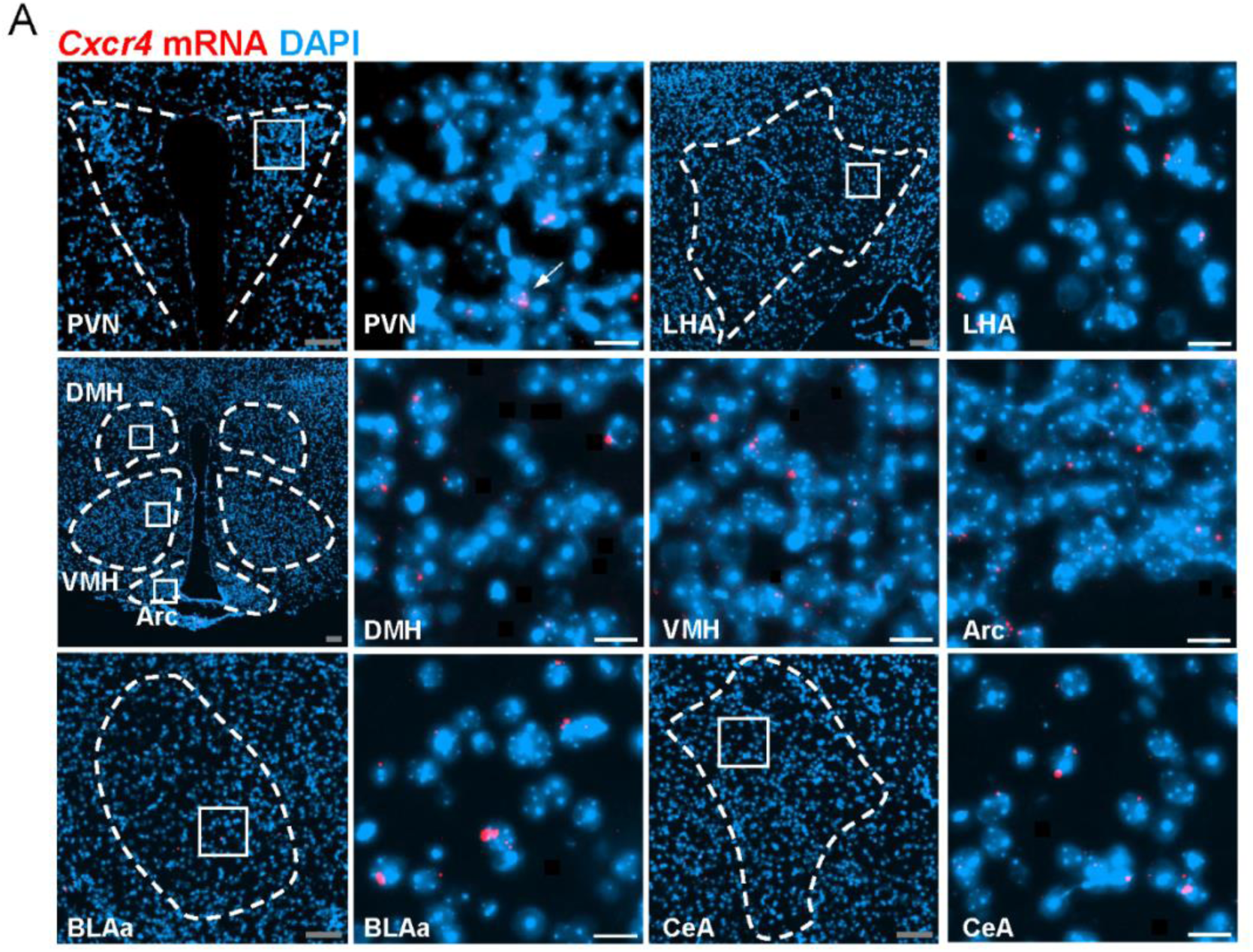
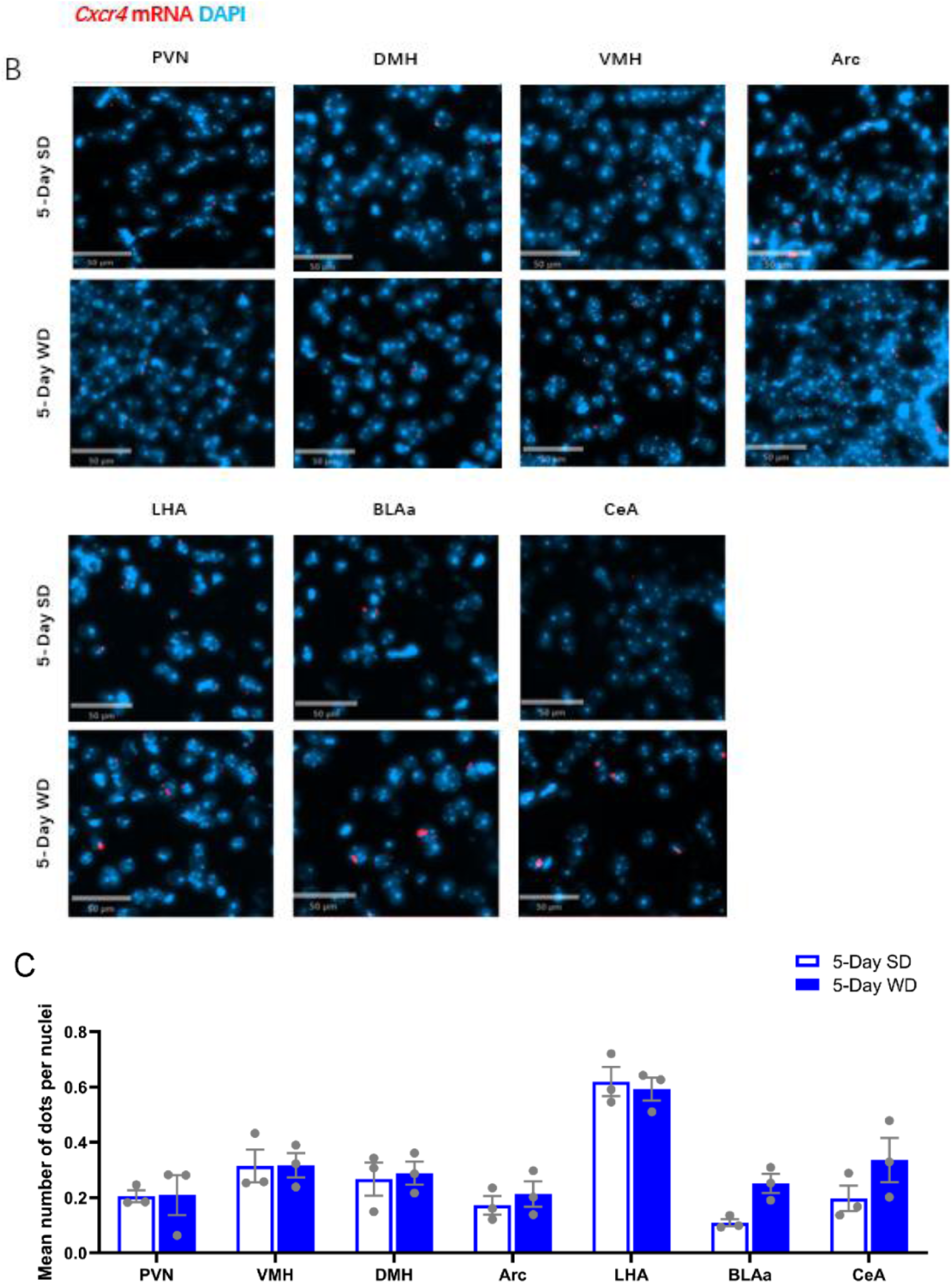
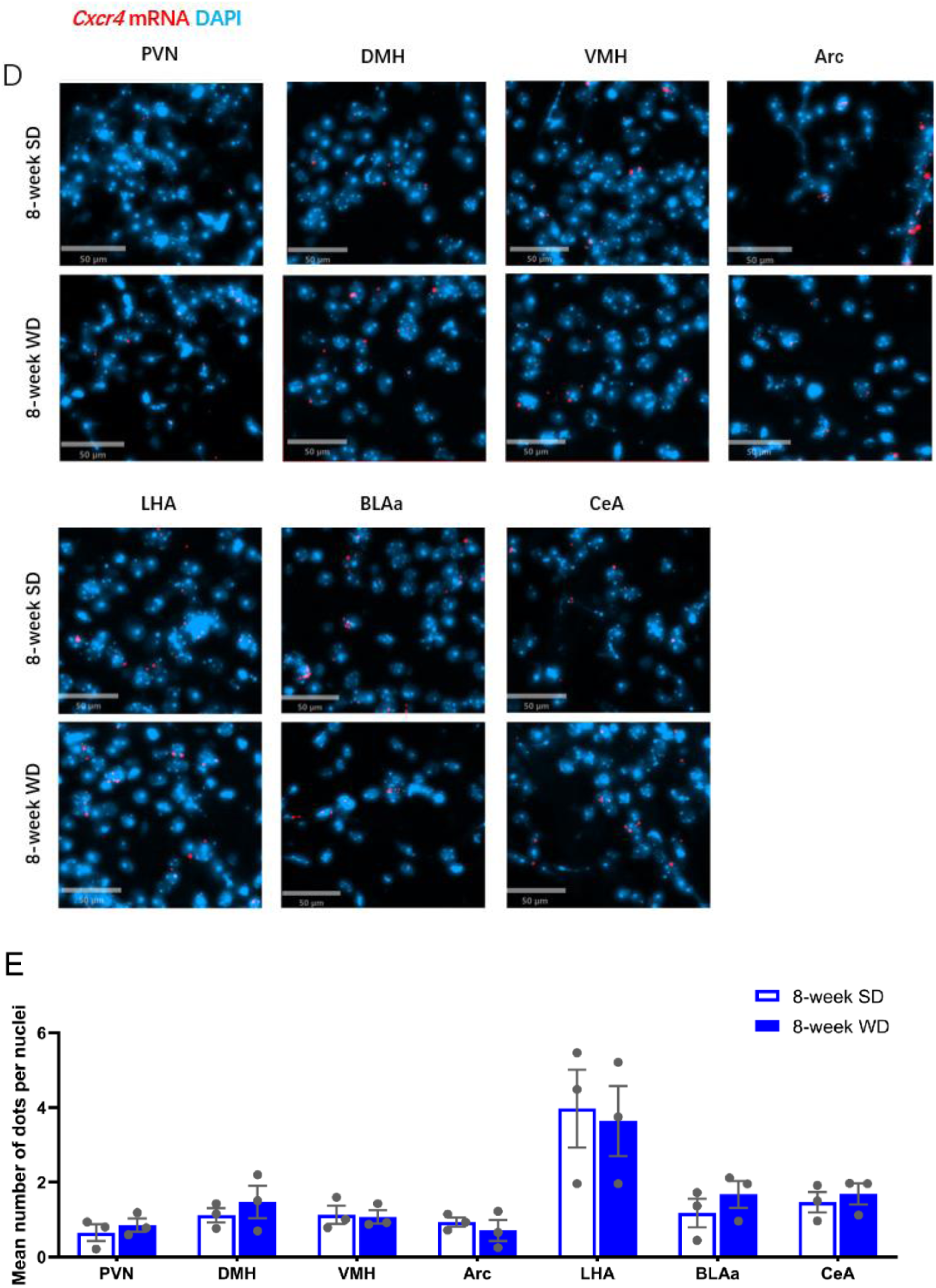
*Cxcr4* expression in hypothalamic and amygdala subregions following short-term (5-day) and prolonged (8-week) WD exposure in male *Ldlr^⁻^*^/^*^⁻^* mice. (A) RNAscope in situ hybridization detection of *Cxcr4* mRNA (red) in DAPI-stained (blue) in the PVN, DMH, VMH, Arc, LHA, BLAa, CeA. Scale bar: 20 µm. (B) RNAscope detection of *Cxcr4* mRNA in brain regions including the PVN, DMH, VMH, LHA, Arc, BLAa and CeA, following a 5-day SD or WD exposure. Representative images show *Cxcr4* mRNA (red dots) with DAPI (blue); scale bar = 20 µm. (C) Corresponding quantification represents the mean number of *Cxcr4* mRNA dots per cell across the indicated brain regions (PVN, DMH, VMH, LHA, Arc, BLAa and CeA) following a 5-day SD or WD exposure. Data are presented mean ± SEM, n = 3 mice (3 sections/mouse); Mann–Whitney U test. (D) RNAscope detection of *Cxcr4* mRNA in brain regions including the PVN, DMH, VMH, LHA, Arc, BLAa and CeA, following an 8-week SD or WD exposure. Representative images show *Cxcr4* mRNA (red dots) with DAPI (blue); scale bar = 20 µm. (E) Corresponding quantification represents the mean number of *Cxcr4* mRNA dots per cell across the indicated brain regions (PVN, DMH, VMH, LHA, Arc, BLAa and CeA) following an 8-week SD or WD exposure. Data are presented mean ± SEM, n = 3 mice (3 sections/mouse); Mann–Whitney U test.

Thaler et al. demonstrated that diet-induced obesity is accompanied with an early hypothalamic inflammatory response, characterized by a transient upregulation of proinflammatory gene expression within the first week of high-fat diet (HFD) exposure [26]. Based on this finding, we investigated whether WD–induced atherogenesis in male *Ldlr^⁻/⁻^* mice is accompanied by a similar transient alteration in *Cxcr4* gene expression, with a specific focus on hypothalamic subregions (PVN, DMH, VMH, Arc, LHA) as well as amygdala subregions (BLAa, CeA). Based on previous studies [27], we selected 5 days of WD feeding to capture potential early-phase changes and 8 weeks WD feeding to evaluate long-term adaptations in the expression of *Cxcr4*. RNAscope analysis revealed no significant differences in *Cxcr4* mRNA expression in the PVN, DMH, VMH, LHA, Arc, BLAa and CeA after 5 days of WD exposure compared to SD (Figure 4B, C). Similarly, prolonged 8-week WD feeding did not result in significant alterations in *Cxcr4* mRNA expression in these regions (Figure 4D, E). Together, our analyses revealed no transient or long-term changes in the consistently low expression of *Cxcr4* in hypothalamic and amygdala subregions during WD-induced atherogenesis of male *Ldlr^⁻/⁻^* mice.

### PVN *Mif* gene expression shows transient upregulation in response to WD feeding in male *Ldlr^⁻^*^/^*^⁻^* mice

MIF, a non-cognate ligand of the chemokine receptor CXCR4, activates CXCR4 signaling and promotes the recruitment of atherogenic leukocytes, thereby exacerbating inflammation within atherosclerotic lesions [11], is also expressed in brain [28, 29] and has previously been implicated in neuroendocrine-immune crosstalk [30, 31]. We again performed RNAscope to map *Mif* expression in brain regions involved in neuroimmune regulation of atherosclerosis, focusing on hypothalamic subregions (PVN, DMH, VMH, Arc, LHA) and amygdala subregions (BLAa, CeA). RNAscope results showed highly abundant *Mif* transcripts, visualized as green puncta, in the PVN, DMH, VMH, Arc, LHA, BLAa, and CeA (Figure 5A). Following the experimental design applied in our CXCR4 studies, we analyzed *Mif* mRNA expression by RNAscope after 5 days and 8 weeks of WD exposure in male *Ldlr^⁻/⁻^* mice. A significant increase in *Mif* mRNA expression was observed exclusively in the PVN after 5 days of WD feeding, whereas no significant changes were detected in other hypothalamic or amygdala subregions compared with SD controls (Figure 5B, C). After 8 weeks of WD exposure, *Mif* mRNA expression in the PVN returned to baseline, with no alterations in other brain regions (Figure 5D, E). These findings indicate that *Mif* mRNA expression in the PVN is transiently upregulated during early WD exposure, potentially contributing to neuroimmune signaling events in the initial phase of WD-induced atherogenesis of male *Ldlr*^⁻/⁻^ mice.

**Figure 5.**
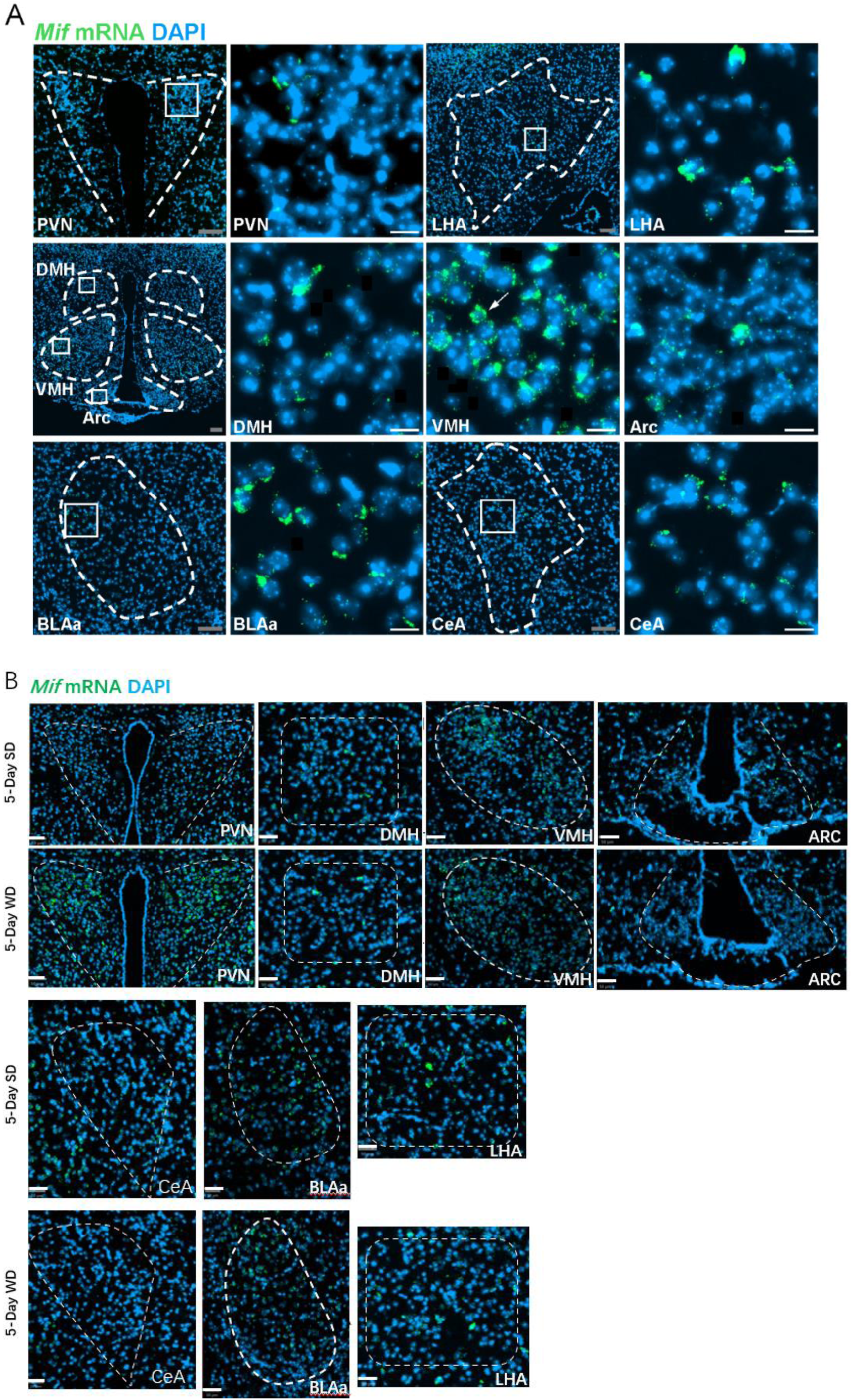
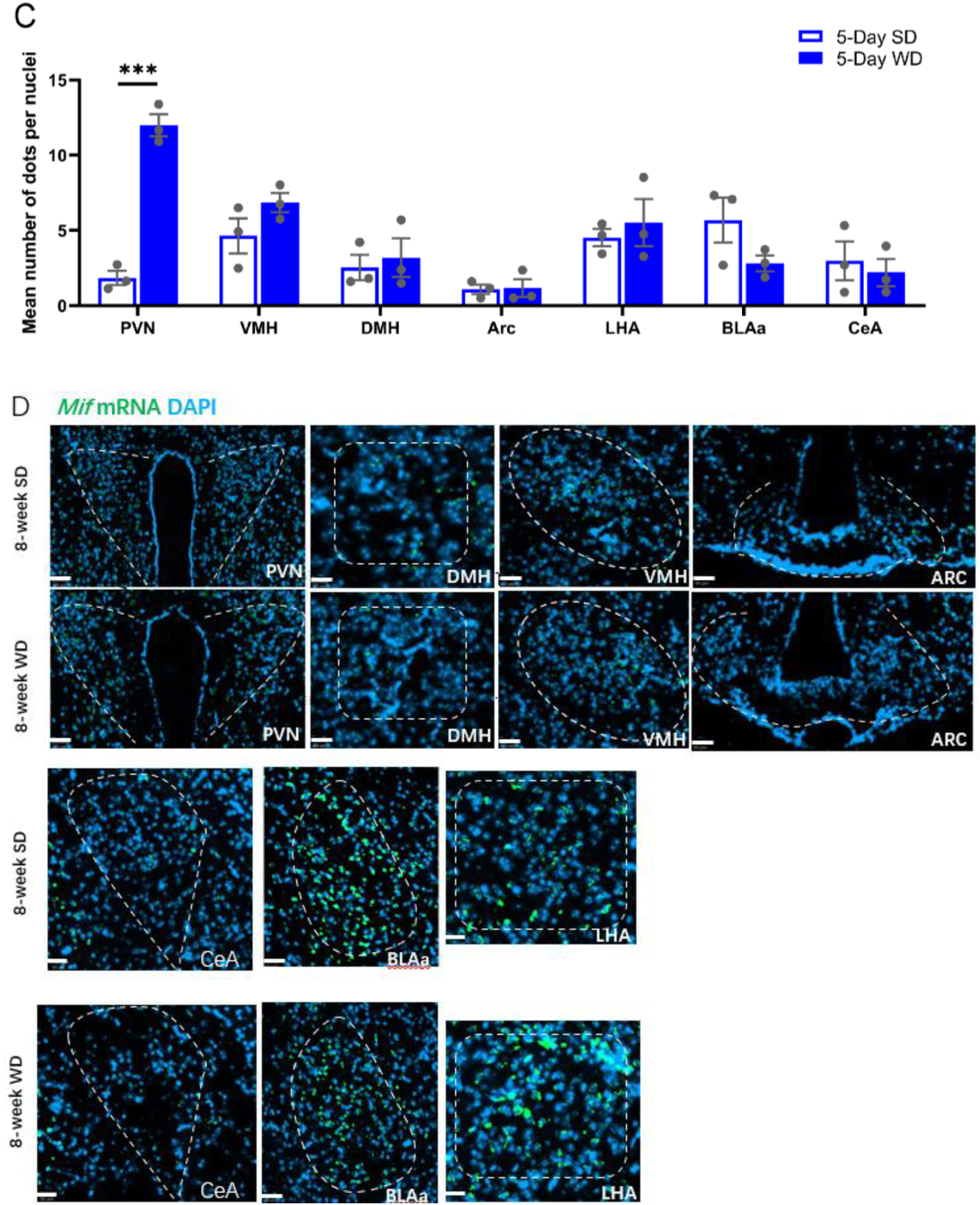
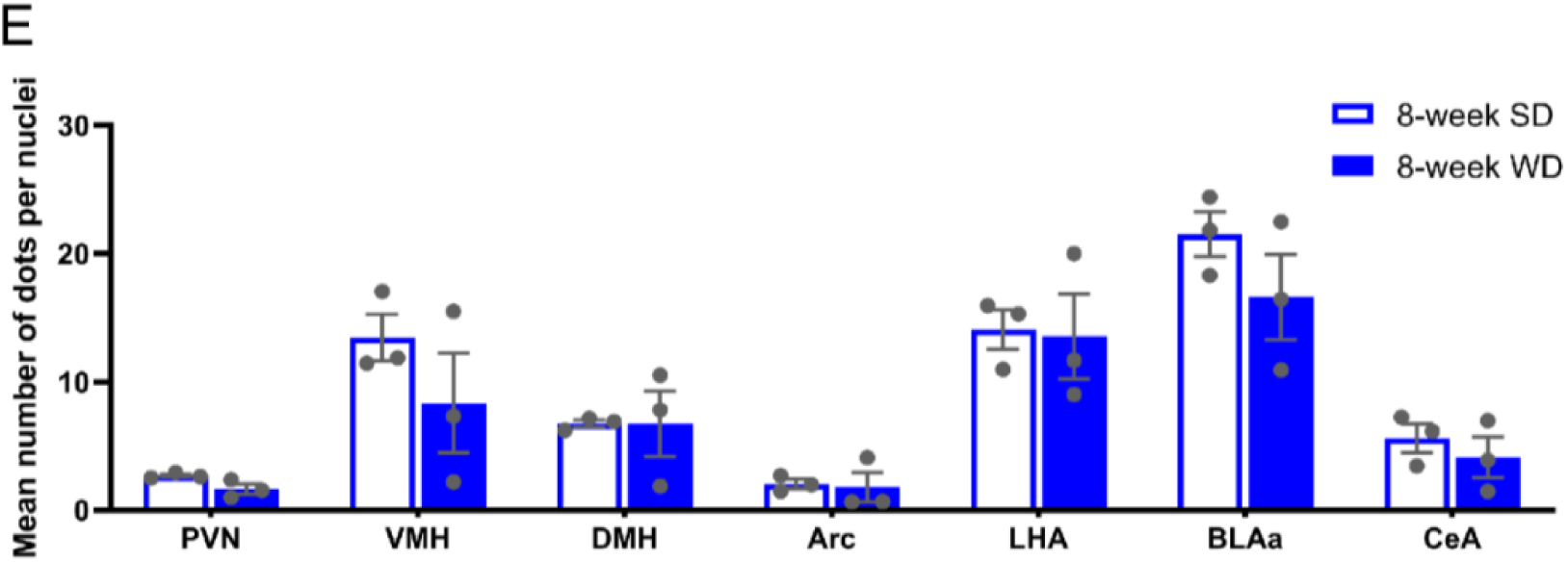
*Mif expression* in hypothalamic and amygdala subregions following short-term (5-day) and prolonged (8-week) WD exposure in male *Ldlr^⁻^*^/^*^⁻^* mice. (A) RNAscope in situ hybridization detection of *Mif* mRNA (green) in DAPI-stained (blue) in the PVN, DMH, VMH, Arc, LHA, BLAa, CeA. Scale bar: 50 µm. White arrows indicate representative *Mif* mRNA–positive cells. scale bar = 20 µm. (B) RNAscope detection of *Mif* mRNA in brain regions including the PVN, DMH, VMH, LHA, Arc, BLAa and CeA, following a 5-day SD or WD exposure. Representative images show *Mif* mRNA (green dots) with DAPI (blue); scale bar = 20 µm. (C) Corresponding quantification represents the mean number of *Mif* mRNA dots per cell across the indicated brain regions (PVN, DMH, VMH, LHA, Arc, BLAa and CeA) following a 5-day SD or WD exposure. Data are presented mean ± SEM. n = 3 mice, 4 coronal sections per mouse (Mann-Whitney U test, ***, *p* < 0.05). (D) RNAscope detection of *Mif* mRNA in brain regions including the PVN, DMH, VMH, LHA, Arc, BLAa and CeA, following an 8-week SD or WD exposure. Representative images show *Mif* mRNA (green dots) with DAPI (blue); scale bar = 20 µm. (E) Corresponding quantification represents the mean number of *Mif* mRNA dots per cell across the indicated brain regions (PVN, DMH, VMH, Arc, LHA, BLAa and CeA) following an 8-week SD or WD exposure. Data are presented mean ± SEM. n = 3 mice, 3 coronal sections per mouse (Mann-Whitney U test).

### Exploratory evidence suggests that short-term WD may activate MIF–CXCR4 signaling in the PVN of male *Ldlr^⁻/⁻^* mice

Based on our RNAscope results showing that *Mif* expression in the PVN is transiently upregulated in response to WD feeding in male *Ldlr*^⁻/⁻^ mice, we next performed 10x Visium spatial transcriptomic sequencing on coronal brain sections containing the PVN regions to investigate whether and how short-term WD exposure activates MIF–CXCR4 signaling and its downstream pathways. After quality control, 11,570 spatial spots were retained for analysis from two experimental conditions: one mouse fed a SD for 5 days (Sample_SD; 5,823 spatial spots) and one mouse fed a WD for 5 days (Sample_WD; 5,747 spatial spots). Unsupervised clustering of all these spatial spots, performed by PCA-based dimensionality reduction followed by Louvain graph-based clustering (resolution = 1.0), identified 29 distinct transcriptomic clusters. Visualization in UMAP space revealed clear separation of these clusters (Figure 6A), and their spatial distributions across coronal brain sections are shown in Figure 6B.

**Figure 6.**
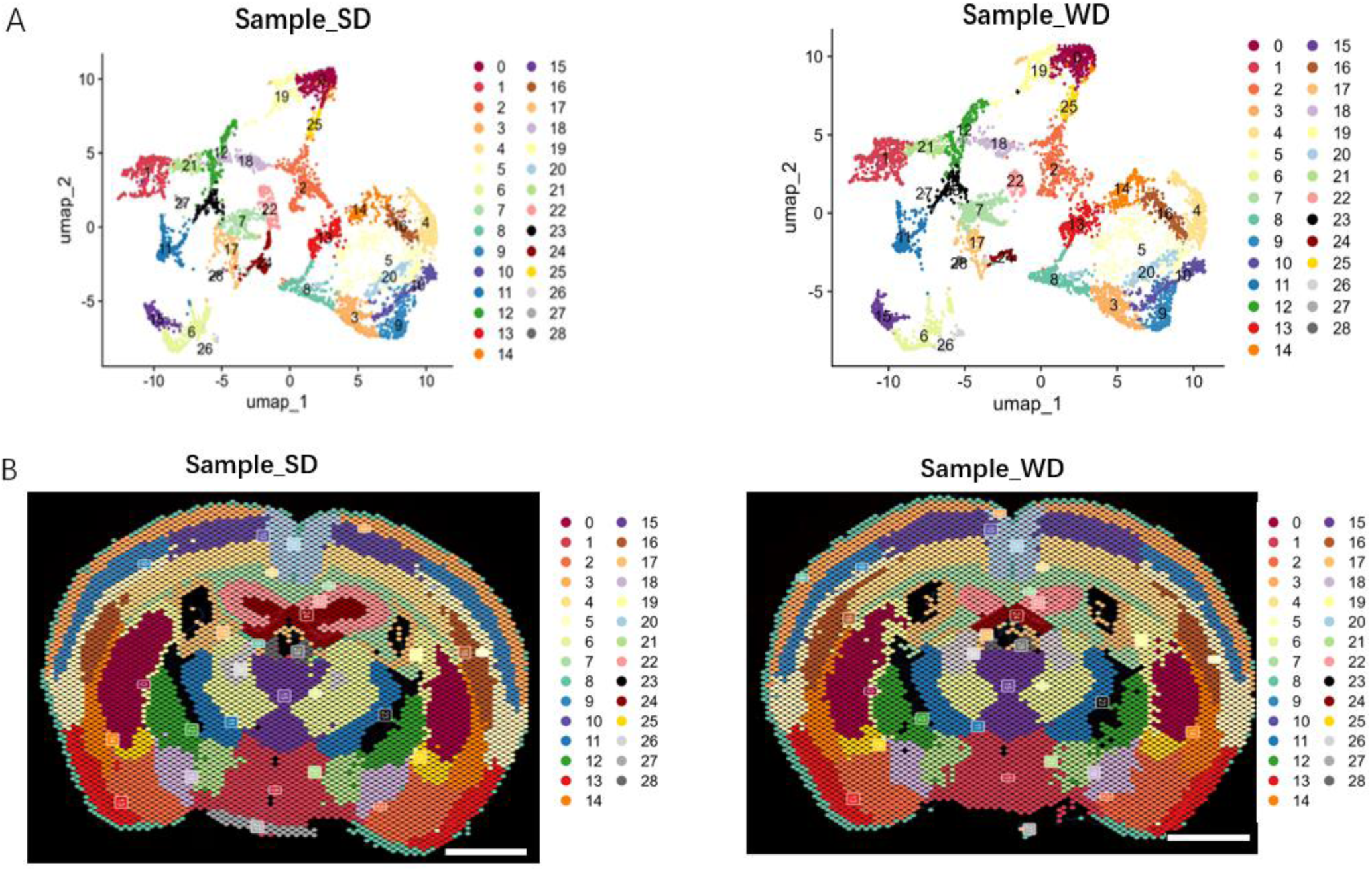
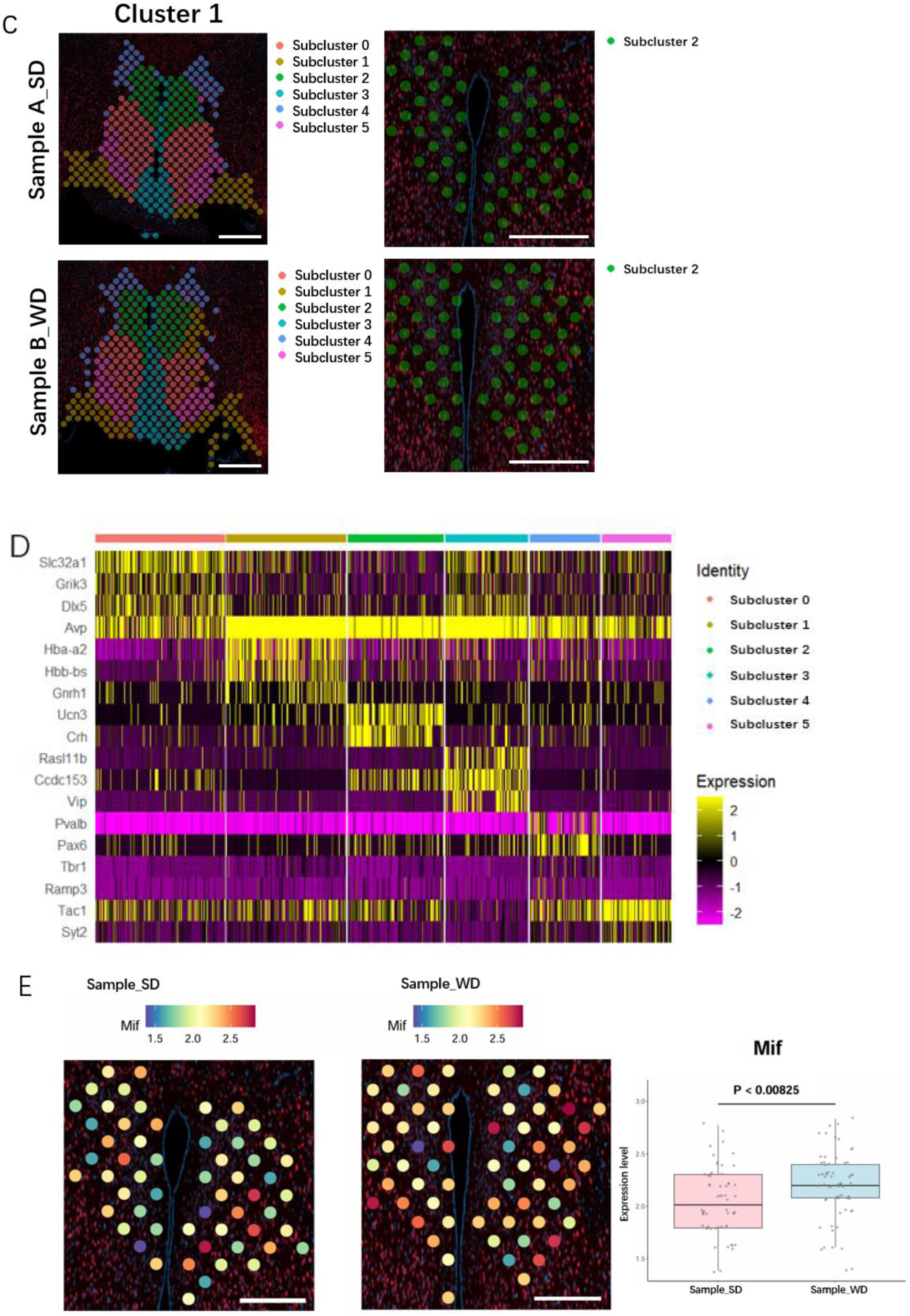
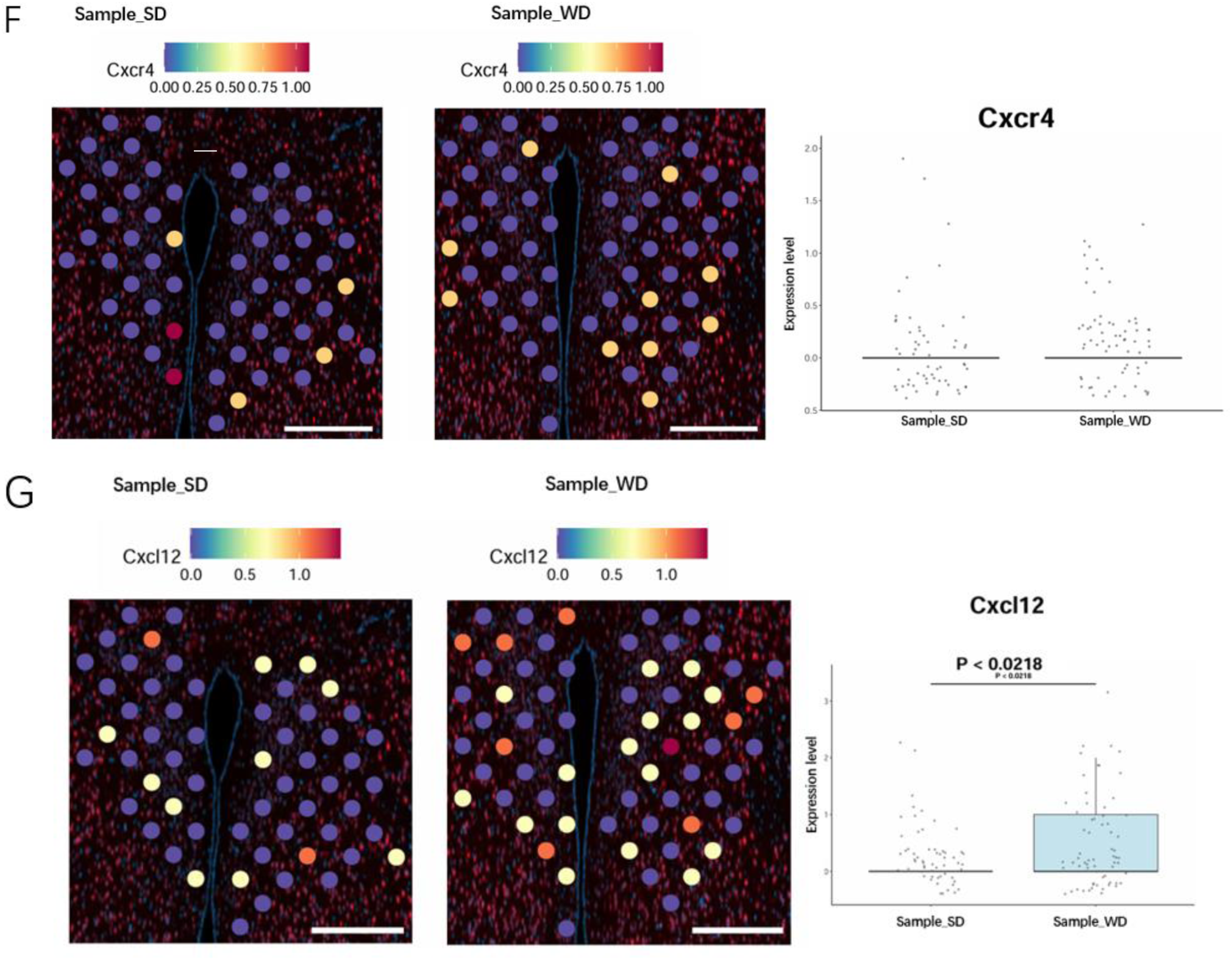
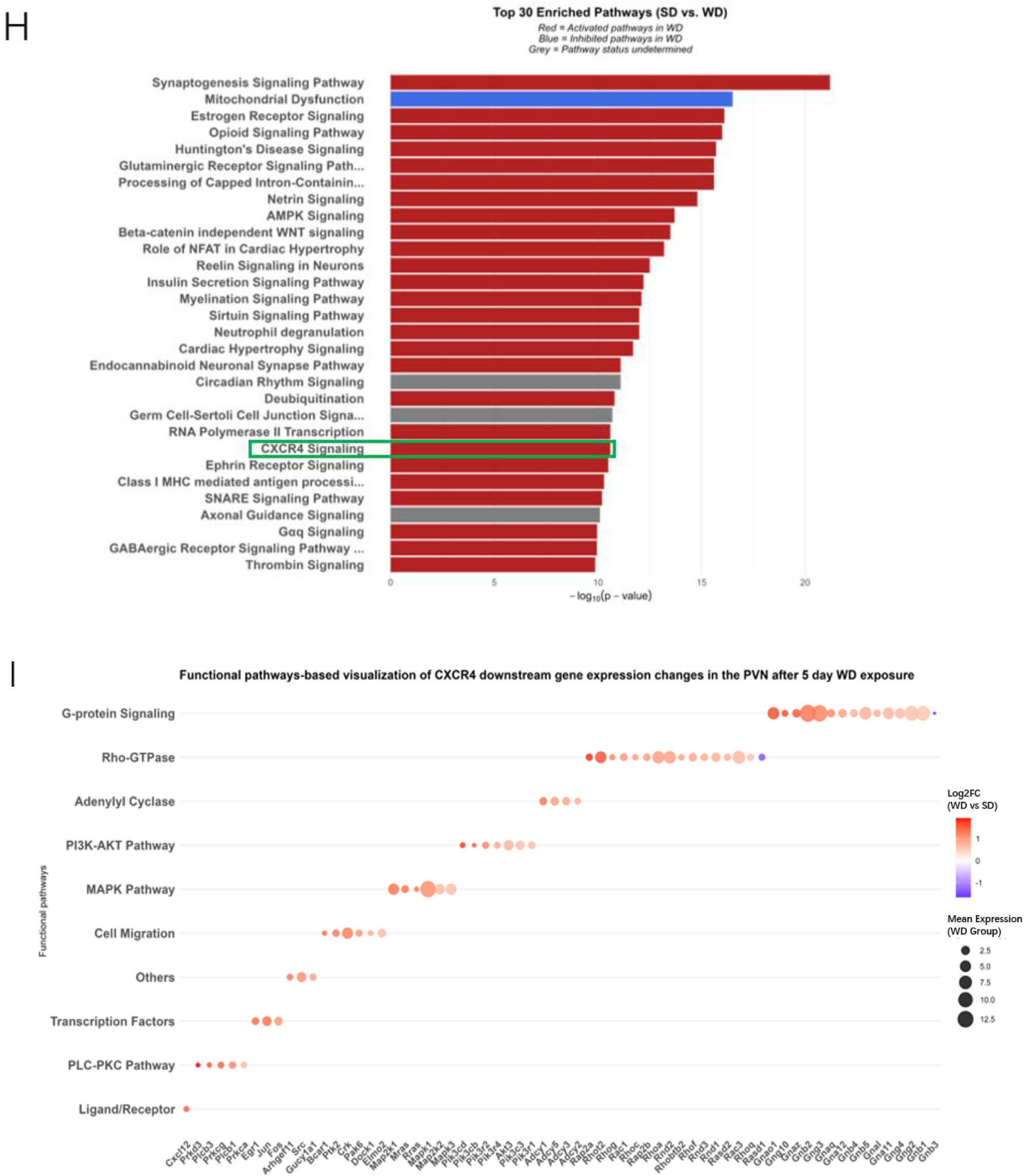
Exploratory spatial transcriptomics identifies a PVN-enriched cluster and reveals short-term WD-induced activation of the MIF–CXCL12/CXCR4 signaling pathway in PVN of male *Ldlr*^⁻/⁻^ mice. (A) UMAP plots showing the distribution of 29 clusters in Sample A_SD and Sample B_WD. Different clusters are color-coded. (B) Spatial plots showing the distribution of 29 clusters in coronal brain sections of Sample A_SD and Sample B_WD. Each cluster is color-coded consistently with the UMAP plot. Scale bar: 100 µm. (C) Spatial plots showing the distribution of six subclusters identified from Cluster 1 (left) and the Subcluster 2 (the PVN-enriched cluster, right) in Sample_SD and Sample_WD. Scale bar, 50 µm. (D) Heatmap showing the top five marker genes for each of the six subclusters within Cluster 1. Subclusters are indicated by distinct colors, and normalized gene expression is represented from low (purple/black) to high (yellow). (E–G) Spatial plots (left panel) showing the *Mif* (E), *Cxcr4* (F), and *Cxcl12* (G) expression within the PVN-enriched cluster in Sample_SD and Sample_WD. Boxplots (right panel) showing the corresponding expression levels in the PVN-enriched cluster between Sample A_SD (pink) and Sample B_WD (light blue). Boxes represent the median and interquartile range, with individual dots representing normalized expression values from single spatial spots. Statistical comparisons between diet groups were performed using the Wilcoxon rank-sum test. Scale bar, 50 µm. (H) Top 30 significantly enriched canonical pathways identified by IPA of DEGs between Sample_SD and Sample_WD in the PVN-enriched cluster. Pathways are ranked by –log_10_(*p*). Red = activated pathways in WD; blue = inhibited pathways in WD; grey = pathways with undetermined activation status. The CXCR4 signaling pathway is highlighted in a green rectangle. (I) Bubble plot of genes annotated by IPA as members of the canonical CXCR4 signaling pathway, categorized into functional modules (G-protein signaling, adenylyl cyclase, PLC–PKC, PI3K–AKT, MAPK, Rho-GTPase, cell migration, transcription factors, ligand/receptor, and others). Bubble size represents mean normalized expression in the WD group, and bubble color represents log_2_ fold-change (WD vs SD) on a blue–white–red scale centered at zero. Visualization of downstream CXCR4 signaling pathway-associated genes in the PVN following short-term WD exposure, categorized by functional pathways. Dot size represents the mean normalized expression level of each gene, and color intensity indicates the log₂ fold-change (WD vs. SD). Functional modules include canonical CXCR4 signaling effectors such as G-protein signaling, Rho-GTPase, Adenylyl cyclase, PI3K–AKT pathway, MAPK pathway, Cell migration, Transcription factors, PLC–PKC pathway.

Among these clusters, Cluster 1 anatomically encompassed the PVN region of interest. A total of 725 spatial spots from Cluster 1 (357 spots from Sample_SD and 368 spots from Sample_WD) were further divided into six subclusters by PCA-based dimensionality reduction followed by Louvain graph-based clustering (resolution = 0.5). Notably, Subcluster 2 showed the closest spatial mapping with the PVN (Figure 6C). Marker gene analysis was then performed across all six subclusters within Cluster 1 (725 spots pooled from both diets) to identify subcluster-specific marker genes. Heatmap visualization of the top five markers per subcluster showed that *Avp, Crh* and *Ucn3*, canonical markers of PVN neurons, were robustly enriched in Subcluster 2 (Figure 6D) [32–34]. While *Avp* was strongly expressed in Subcluster 2, it was also detected in other subclusters, consistent with its known distribution not only in PVN neurons but also in adjacent hypothalamic nuclei [35]. The co-enrichment of the PVN marker genes (*Crh*, *Ucn3*, and *Avp*) within Subcluster 2, together with its spatial overlap with the PVN, provides strong evidence for identifying this Subcluster 2 as PVN-enriched cluster. 5-Day WD feeding significantly increased *Mif* expression in the PVN-enriched cluster, whereas *Cxcr4* expression remained unchanged (Figure 6E, F), consistent with our RNAscope findings. Notably, the canonical CXCR4 ligand *Cxcl12* was also upregulated after 5-Day WD feeding (Figure 6G). IPA on DEGs between Sample_SD and Sample_WD in the PVN-enriched cluster showed significant activation of the CXCR4 signaling pathway (Figure 6H). Together, these results suggest that short-term WD exposure activates CXCR4 signaling through dual-ligand induction (*Mif* and *Cxcl12*) without detectable changes in receptor (*Cxcr4*) expression.

To specifically characterize CXCR4 signaling pathway activation, genes annotated by IPA as members of the canonical CXCR4 signaling pathway were visualized and categorized into functional pathways (Figure 6I). These results revealed CXCR4 signaling in the PVN via dual-ligand (MIF and CXCL12) activation was accompanied by the enrichment of key downstream functional pathways—including G-protein signaling, Rho-GTPase, adenylyl cyclase, PI3K–AKT pathway, MAPK pathway, cell migration, transcription factors, PLC–PKC pathway—all of which are canonical effectors of CXCR4 activation.

## Discussion

Herein we identify a previously unrecognized role of neural CXCR4 in modulating atherosclerotic plaque formation in the cardiometabolic *Ldlr^⁻/⁻^* mouse model, whereby neural-specific *Cxcr4* deletion significantly reduces lesion burden in the aortic arch and whole aorta. Notably, this protective effect was region-specific, as lesion size at the aortic root remained unchanged, indicating a region-specific effect of neural Cxcr4 deletion on atherogenesis. Considering the CXCR4/MIF axis as key pro-atherogenic axis [11, 12], this observation is in line with previous data showing region-specific reduction of plaques in *Mif*-deficient mice on an atherogenic *Apoe*^-/-^ background, in which protection was seen in aortic arch and abdominal aorta, but not root [36, 37].

Our observation that *Mif* gene expression in the PVN increases transiently is consistent with previous evidence that short-term high-fat diet feeding induces transient hypothalamic inflammation and proinflammatory gene expression [26], thereby priming the PVN for subsequent neuroinflammatory and systemic cardiometabolic pathology. By examining short-term (5-day) WD exposure in *Ldlr^⁻/⁻^* mice, we were able to distinguish this early response from secondary effects of weight gain and advanced metabolic syndrome. We found that short-term WD feeding did not change *Cxcr4* mRNA levels in the PVN, yet our exploratory spatial transcriptomics data showed suggestive trends that the CXCR4 pathway may be activated, as predicted by IPA and supported by downstream gene enrichment. This pattern reflects a classical GPCR paradigm, in which ligand availability, rather than receptor abundance, dictates signaling strength through cascades such as G proteins, PI3K–AKT, MAPK, and Rho GTPases [38, 39]. The concurrent upregulation of the non-canonical ligand *Mif* and potentially also of the canonical ligand *Cxcl12* suggests that early dietary stress may engage CXCR4 signaling via a ligand-driven mechanism. Although our findings are in contrast with studies in rats that reported increased CXCR4 expression in the PVN after 5 days of HFD [27], these discrepancies could reflect species-specific regulation (mouse vs. rat), differences in genetic background (wild type vs. *Ldlr⁻^/^⁻*), variations in diet composition, or differences in the timing and sensitivity of detection methods. Another possibility is that CXCR4 regulation under dietary stress is context-dependent: while some studies have reported increased receptor expression, others have described stable receptor levels with ligand-driven activation as the predominant mechanism [40], consistent with our observations.

Our findings provide insights into how nutrient-induced CXCR4 signaling may contribute to neuroimmune adaptations in the PVN during the early phase of atherogenesis. Our explorative evidence suggests that short-term WD feeding may activate canonical CXCR4 downstream cascades, including PI3K–AKT and MAPK, which are classically associated with cell survival, proliferation, and migration [41] and are also implicated in synaptic plasticity and inflammatory responses in neurons [42, 43].Of note, the MIF/CXCR4 axis has previously been associated with the activation of the PI3K/AKT pathway [44, 45], supporting our conclusion. In addition, we observed enrichment of Rho-GTPase signaling, a branch frequently engaged downstream of CXCR4 [46]. Rho GTPases are well known to regulate actin cytoskeleton remodeling and membrane morphology [47, 48] and to influence cell migration and adhesion in CXCR4 mediated signaling [46]. This enrichment suggests that CXCR4 activation under dietary stress could drive structural plasticity of PVN neurons and/or promote glial activation and migration, processes that may constitute the substrate of early hypothalamic inflammation and dysfunction. Notably, recent studies have identified hypothalamic glial remodeling as an early hallmark of diet-induced metabolic disease [49–51], supporting our interpretation. Importantly, our exploratory spatial transcriptomics analysis supports WD-induced *Mif* expression localized to the PVN, consistent with our RNAscope findings and distinct from other hypothalamic regions. Given the PVN’s central role in autonomic and cardiovascular regulation, the CXCR4 chemokine axis identified here may represent a previously unrecognized neuroimmune conduit modulating vascular inflammation and the progression of atherosclerosis.

Limitations of the present study include (i) use of *Nestin-Cre*, which can affect progenitors/glia and complicate cell-type specificity interpretation [52, 53], and (ii) temporal sampling at only two time points. Future investigations should dissect cell-type-specific CXCR4 signaling in PVN and test ligand-selective perturbations (MIF vs. CXCL12) to elucidate their contributions to autonomic regulation, peripheral inflammation, and plaque formation.

## Conclusions

This study identifies neural CXCR4 as a component of CNS-mediated modulation of atherosclerotic plaque formation, exerting its effects independent of systemic metabolic or inflammatory changes. Short-term Western diet exposure induced gene expression of the CXCR4 ligand MIF in the PVN. Combined, these findings establish a connection between CNS CXCR4 function and vascular disease, highlighting the role of brain–immune–vascular interactions in cardiometabolic risk and pointing toward potential avenues for CNS-targeted intervention strategies.

## List of Abbreviations

CNS: Central nervous system
CXCR4: C-X-C motif chemokine receptor 4
Ldlr^⁻/⁻^: Low-density lipoprotein receptor–deficient
WD: Western diet
MIF: Macrophage migration inhibitory factor
PVN: Paraventricular nucleus
GPCR: G protein–coupled receptor
Apoe^⁻/⁻^: Apolipoprotein E–deficient
IF: Immunofluorescence
DAPI: 4′,6-diamidino-2-phenylindole
DEGs: Differentially expressed genes
Crh: Corticotropin-releasing hormone
Ucn3: Urocortin 3
Avp: Arginine vasopressin
IPA: Ingenuity Pathway Analysis
ARC: Arcuate nucleus
VLDL: Very low density lipoprotein
LDL: Low density lipoprotein
HDL: High density lipoprotein
WBC: White blood cells
RBC: Red blood cells
PLT: Platelets
DMH: Dorsomedial hypothalamus
VMH: Ventromedial hypothalamus
LHA: Lateral hypothalamic area
BLAa: Basolateral amygdala, anterior part
CeA: Central amygdala
HFD: High-fat diet
UMAP: Uniform manifold approximation and projection

## Declarations

### Ethics approval and consent to participate

All procedures were approved by the local Animal Use and Care Committee and the local authorities of Upper Bavaria, Germany in accordance with European and German animal welfare regulations.

### Consent for publication

All authors have declared their consent for this publication.

### Availability of data and materials

The datasets used and/or analysed during the current study are available from the corresponding author upon reasonable request.

### Competing interests

T.D.M. receives research funding from Novo Nordisk, the German Research Foundation (DFG TRR296, TRR152 and GRK 2816/1) and the European Research Council ERC-CoG Trusted no. 101044445, but these funds are unrelated to the here described work. T.D.M. receives research funding by Novo Nordisk and has received speaking fees from Novo Nordisk, Eli Lilly, Boehringer Ingelheim, Merck, AstraZeneca and Mercodia. S.M.H. receives research funding from the German Research Foundation (FOR 5298) that is unrelated to the here described work. J.B. and C.W. are co-inventors of patents covering anti-MIF strategies for inflammatory and cardiovascular diseases. A.K. and J.B. are co-inventors of a patent application covering MIF-binding CXCR4 ectodomain mimics for inflammatory and cardiovascular diseases.

### Funding

Open Access funding enabled and organized by Project DEAL. This work was supported in part by funding S.M.H., T.D.M, J.B., Y.D. and C.W. through the Deutsche Forschungsgemeinschaft (DFG; SFB1123-A01 & A04) and the DZD. We also acknowledge support from DFG grant CRC1123-A3 to J.B. and A.K., by DFG under Germany’s Excellence Strategy within the framework of the Munich Cluster for Systems Neurology (EXC 2145 SyNergy-ID 390857198) to J.B., as well as from the m4 award project SELECKREM 41-6663a/214/12-M4-2110-0005 by the Bavarian Ministery of Economic affairs to J.B. and A.K.

### Authors’ contributions

Y.S, R.T. and A.G. generated, analyzed and interpreted experimental data; A.F. assisted in spatial transcriptomics procedure; Y.S. analyzed spatial transcriptomics data and drafted the manuscript; D.L oversaw spatial transcriptomics sequencing data analysis. Y.D. and C.W. performed aortic plaque formation analysis and co-wrote the article; J.B. and A.K. provided MIF reagents and tissues, contributed to the study concept, and revised the manuscript. T.D.M. and C.G.C. contributed to the study concept and critically reviewed the manuscript.

S.M.H and R.Z.T oversaw the in vivo experiments, interpreted experimental data and co-wrote the article. S.M.H. is the guarantor of this work and, as such, has full access to all the data in the study and takes responsibility for the integrity of the data and the accuracy of the data analysis.

## Acknowledgements

We thank Luisa Mu ller, Laura Sehrer, Emiljia Malogajski, Cynthia Striese, Sebastian Cucuruz, Markus Brielmeier at HMGU for excellent assistance with mouse experiments. We thank Ulrike Buchholz, Christina Koupourtidou and Monica Tost at HMGU for excellent technical assistance with spatial transcriptomic wet lab experiments. We thank Ophe lia Le Thuc at HMGU for excellent technical assistance with RNAscope experiment.

## Declaration of AI-assisted technologies in the writing process

We used ChatGPT (model: GPT-5) to improve readability and English language of the manuscript. After using this tool, the authors reviewed and edited the content as needed under strict human oversight and took full responsibility for the content of the publication.

## Supplementary figures

**Supplementary Figure A.**
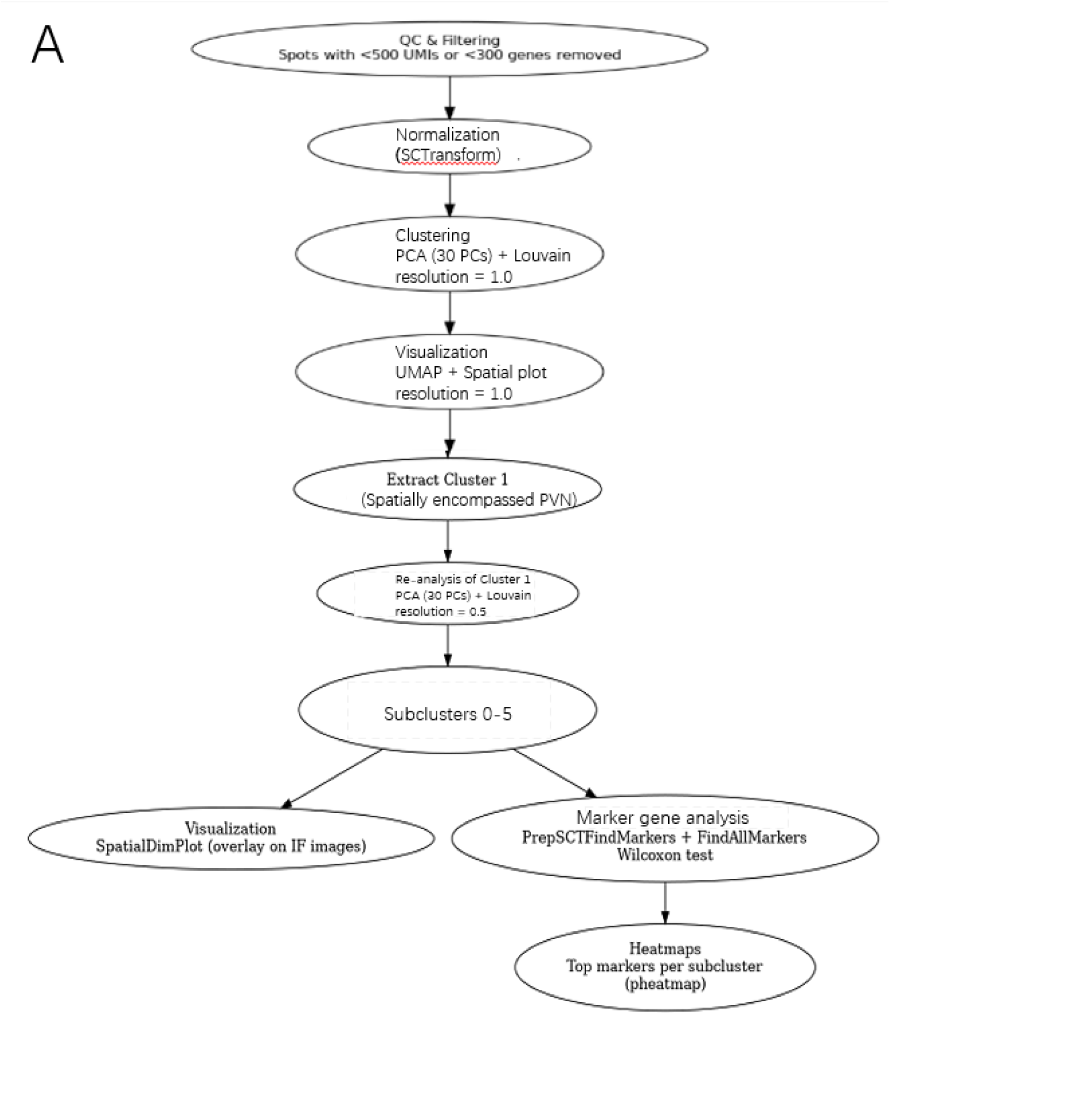
Workflow to identify PVN-enriched cluster.

